# High-throughput behavioral screen in *C. elegans* reveals novel Parkinson’s disease drug candidates

**DOI:** 10.1101/2020.02.20.958751

**Authors:** Salman Sohrabi, Danielle E. Mor, Rachel Kaletsky, William Keyes, Coleen T. Murphy

## Abstract

We recently linked branched-chain amino acid transferase 1 (*BCAT1*) with the movement disorder Parkinson’s disease (PD), and found that reduction of *C. elegans bcat-1* causes abnormal spasm-like ‘curling’ behavior with age. Here, we report the development of a high-throughput automated curling assay and its application to the discovery of new potential PD therapeutics. Four FDA-approved drugs were identified as candidates for late-in-life intervention, with metformin showing the greatest promise for repurposing to PD.

---

Parkinson’s disease (PD) is the most common neurodegenerative movement disorder worldwide^1^. PD is characterized by loss of dopaminergic neurons in the substantia nigra, and formation of abnormal protein aggregates containing α-synuclein^2, 3^. Currently, there is no cure for PD and treatment options only mitigate symptoms without modifying the course of the disease^4^. Using *diseaseQUEST*, our recently developed tissue-network approach to identifying and testing new candidates for human disease genes, we discovered a novel link between branched-chain amino acid transferase 1 (*BCAT1*) and PD^5^. We found that *BCAT1* expression is decreased in PD patient substantia nigra, and reduction of *bcat-1* in *C. elegans* promotes dopaminergic neurodegeneration^5^. Moreover, RNAi-mediated knockdown of *bcat-1* in *C. elegans* causes age-dependent spasm-like ‘curling’ behavior, serving as a new model for PD motor symptoms^5^.

The ability to perform large-scale screening for modifiers of Parkinson’s-like curling dysfunction in *C. elegans* may uncover novel disease mechanisms and identify potential therapeutics. However, manual recording and quantification of curling in a liquid thrashing assay^5^ is labor-intensive and time-consuming, limiting its throughput. Worm tracking software packages have been developed to expedite the analysis of *C. elegans* locomotion^6–9^; however these algorithms typically use videos to track individual worms in consecutive frames, and as a result require analysis of large amounts of image data. In addition, tracking *C. elegans* swimming behavior using morphological analysis requires even higher computing power since worms move faster in a liquid environment compared to solid media^10^. Most importantly, these packages do not effectively track worms in a curled posture, and therefore severely underestimate worm curling^5^.

We have now developed a method to quantify *C. elegans* curling behavior in a high-throughput manner using only a series of snapshots rather than videos. Our automated curling assay uses a motorized stage for capturing snapshots, paired with user-friendly software for curling detection and quantification. This simple and reliable workflow enabled us to analyze more than 32,000 worms to investigate the mechanisms by which *bcat-1* reduction is neurotoxic^11^ and to identify new potential treatments for Parkinson’s disease.

To carry out the manual thrashing assay to quantify age-and/or disease-related motor defects in *C. elegans*, ~100 worms per experimental condition are deposited 10-15 at a time into a 10µL drop of M9 buffer on a microscope slide, and 30s videos are recorded. A standard stopwatch is used to measure the percentage of time spent in a curled pose for each individual worm over the span of each 30s video. As an example, to analyze five potential drug candidates and two relevant controls (Figure 1a) using the manual curling assay, it would take a total of 15 hours to capture videos and analyze the data (Figure 1b).

**Figure 1:**
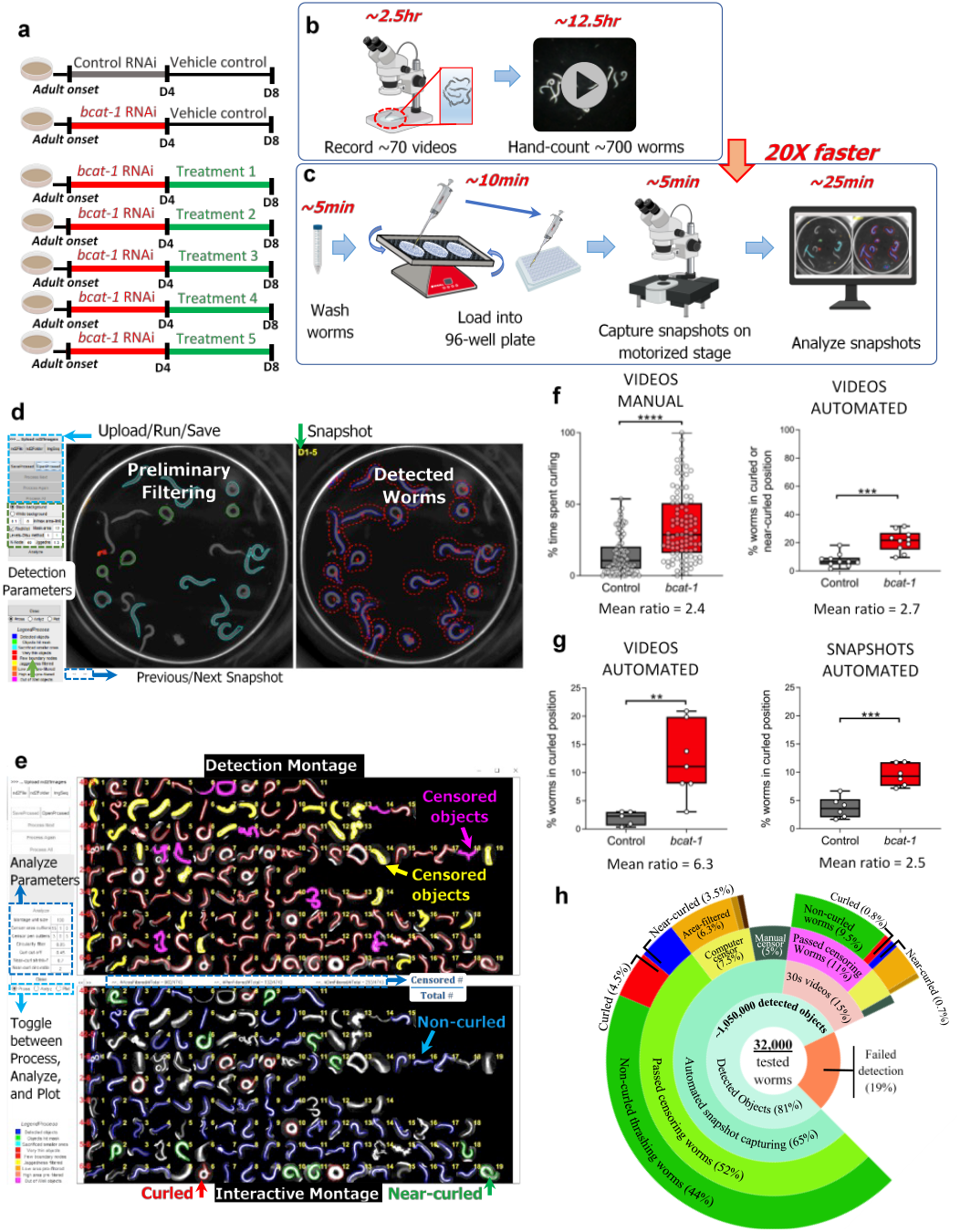
High-throughput automated screening platform for analysis of Parkinson’s-like curling motor dysfunction in *C. elegans.* (a) Experimental design for drug treatments. Neuronal RNAi-sensitive worms were fed *bcat-1* or control RNAi as adults, and on day 4 were transferred to plates with 50-µM drug or vehicle. Curling was measured on day 8. (b) The manual thrashing assay for five drug candidates and two relevant controls requires 15 hours of video recording and hand-counted analysis. (c) The high-throughput automated curling assay can be carried out with only 45min of benchwork for the same or more test conditions. Overall, the automated workflow is 20 times faster than the manual assay. (d) The curling analysis software conducts preliminary filtering and worm detection. Successfully found worms are displayed in blue (right). Scale bar, 2mm. (e) Montage showing successfully detected worms (red) and censored objects (yellow, pink) (top). Worms are categorized as curled (red), near-curled (green), or non-curled (blue) (bottom). (f) Comparison of manual hand-counting (left) versus software analysis (right) for the same set of 30s videos of vehicle-treated worms showed significant curling detected by both methods. The program summed curled and near-curled worms to more closely mimic hand-counting. n = 99 worms for control manual, 98 worms for *bcat-1* manual, 10 videos totaling 99 worms for control automated, 10 videos totaling 98 worms for *bcat-1* automated. (g) Comparison of automated analysis of 30s videos versus snapshots (captured on the same day from two separate aliquots of worms) showed that snapshots are sufficient to detect a significant difference between *bcat-1(RNAi)* and control. n = 5 wells totaling 50 worms for control videos, 7 wells totaling 73 worms for *bcat-1* videos, 6 wells totaling 71 worms for control snapshots, 6 wells totaling 41 worms for *bcat-1* snapshots. (h) Sunburst plot summarizing program performance in detecting and categorizing 32,000 worms. Data are mean±s.e.m. Two-tailed *t*-tests. **p<0.01, ***p<0.001, ****p<0.0001. Box-plots show minimum, 25th percentile, median, 75th percentile, maximum. Mean of bcat-1(RNAi) divided by mean of control RNAi is abbreviated as mean ratio.

In our high-throughput workflow, worms from each treatment group are washed with M9 buffer into 35mm petri dishes secured on an orbital shaker. Pipetting worms from this continuously rocking setup into the wells of a 96-well plate ensures near-uniform distribution of worms across the wells. A motorized microscope stage is programmed to automatically take one snapshot of each filled well until all wells are imaged, and to repeat this process 7 times for a total of 7 snapshots per well. The entire process of washing worms, filling at least 70 wells, and recording images for 7 experimental conditions requires only 20min of benchwork. User intervention in data analysis using our curling quantification software is also minimal (Figure 1c). Overall, our high-throughput automated curling assay can record, process, and analyze experimental data 20 times faster than the manual thrashing assay.

In order to quantify curling, the software imports raw images, outlines worms in each snapshot, and carries out a first-pass filtration step to eliminate overlapping (non-interpretable) worms and debris. Objects filtered in this preliminary stage are color-coded to help the user choose the proper detection parameters (Figure 1d). During the analysis, the program carries out a second and more rigorous round of censoring entangled worms, bacterial chunks, debris, and progeny. Using shape factors such as circularity and convex hull, the program categorizes animals into curled, near-curled, or non-curled subgroups (Figure 1e). Following categorization, the user has the option to manually override automated decisions where the algorithm failed to correctly identify worms. Finally, the percentage of worms that are in a curled or near-curled posture is calculated out of the total number of worms detected.

To test the utility of our automated software and the use of snapshots instead of videos, we conducted several tests using vehicle-treated *bcat-1(RNAi)* and control RNAi worms. First, the same set of 30s videos were manually hand-counted and analyzed using the software, which extracted 30 stills at 1s intervals from each video. The automated analysis was able to detect significantly higher curling in *bcat-1(RNAi)* worms compared to control, as did manual hand-counting (Figure 1f). The ratios of *bcat-1(RNAi)* to control curling levels were similar, 2.4 in the manual assay compared to 2.7 in the automated quantification (Figure 1f). Next, we tested our automated analysis on a set of 30s videos and snapshots that were collected on the same day from two separate aliquots of the same source worms. While the mean curling level of *bcat-1(RNAi)* was 6.3-fold higher than that of control using 30s videos, and only 2.5-fold higher using snapshots, the analysis of snapshots was sufficient to detect a highly significant difference between *bcat-1(RNAi)* and control (Figure 1g), demonstrating the suitability of this workflow for high-throughput screening. Furthermore, we found that using snapshots to quantify the curled position distinguished between *bcat-1(RNAi)* and control groups more effectively than the near-curled position (Figure 2a).

**Figure 2:**
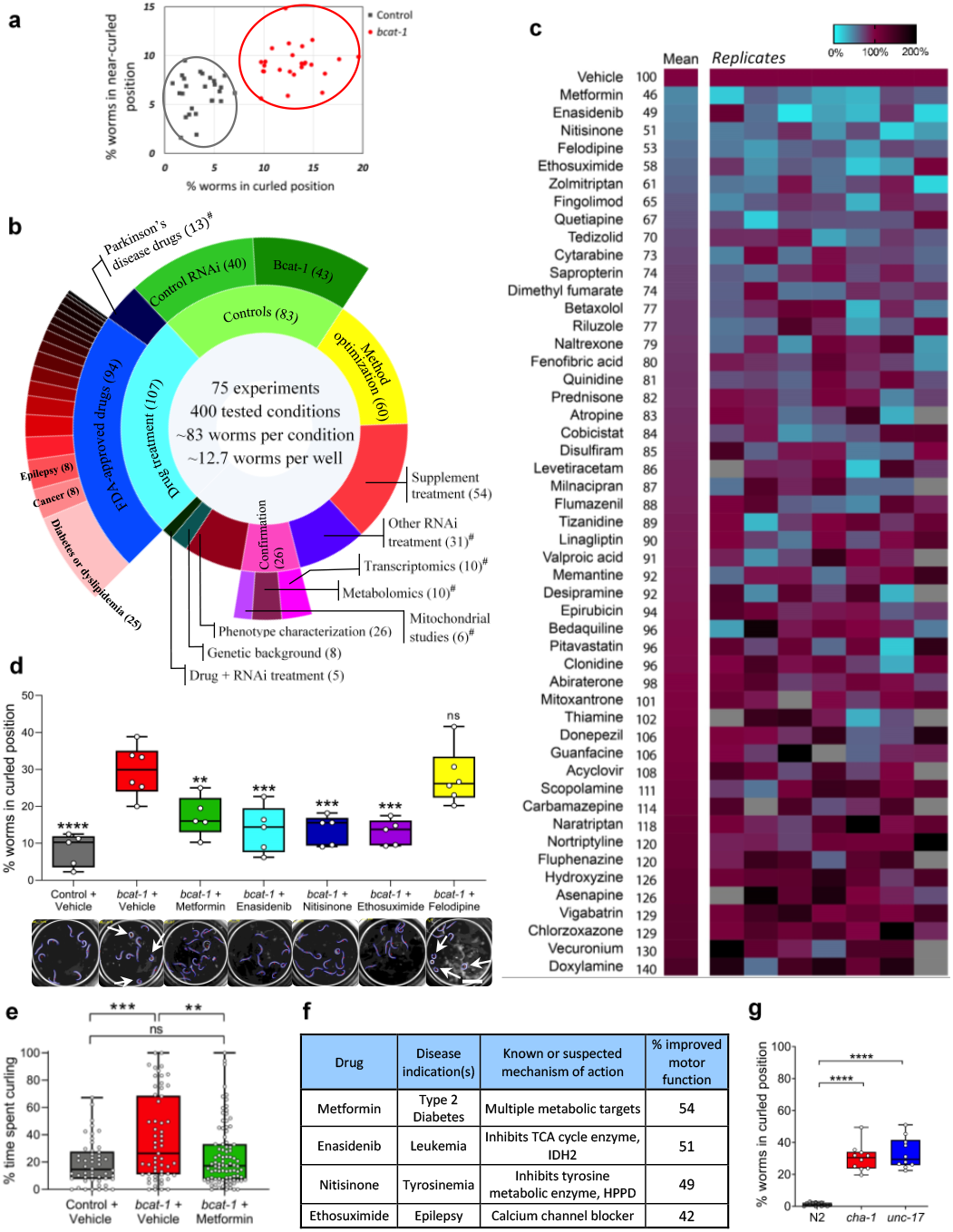
Application of high-throughput curling platform to the discovery of new potential therapeutics for PD. (a) Meta-analysis of curled and near-curled values for vehicle-treated *bcat-1(RNAi)* and control RNAi conditions across n = 26 experiments for control RNAi, 27 experiments for *bcat-1*. Each value represents the mean across all wells in a given experiment. (b) Summary of experimental categories for all 400 conditions tested with number of conditions in parentheses. Results for categories marked with # can be found in our companion study^11^. (c) High-throughput screen of 50 FDA-approved drugs for those that improve motor function in *bcat-1(RNAi)* worms. Vehicle-treated level of curling was set to 100%, and color indicates % curling relative to vehicle. Blank wells are those that had <3 worms and were excluded. (d) Post-screen validation of top five drug candidates, which reduced curling to <60% of vehicle-treated levels. Software analysis of 30s videos confirmed that 4 of the drugs reliably reduce curling. Values were normalized to vehicle-treated *bcat-1(RNAi)* from the same experiment, and statistics are shown for each group compared with *bcat-1(RNAi)* + vehicle. Scale bar, 2mm. n = 5 videos totaling 52 worms for control RNAi, 7 videos totaling 74 worms for *bcat-1(RNAi)* + vehicle, 5 videos totaling 95 worms for metformin, 6 videos totaling 66 worms for enasidenib, 5 videos totaling 83 worms for nitisinone, 5 videos totaling 88 worms for ethosuximide, 6 videos totaling 60 worms for felodipine. One-way ANOVA with Dunnett’s post-hoc. (e) Manual hand-counting served as a final confirmation that the top-performing drug candidate, metformin, significantly improves motor function in the *bcat-1(RNAi*) worm model of PD. n = 50 worms for control vehicle, 57 worms for *bcat-1* vehicle, 91 worms for *bcat-1* metformin. One-way ANOVA with Tukey’s post-hoc. (f) Disease indications and targets of the top 4 drugs that were confirmed to reduce curling. % improved motor function is based on the results from the high-throughput screen. (g) Curling detected in cholinergic mutants *unc-17* and *cha-1* on day 8 indicates that the high-throughput curling assay can be applied beyond the *bcat-1(RNAi)* model to other genetic backgrounds. n = 52 worms for N2, 43 worms for *cha-1*, 57 worms for *unc-17*. One-way ANOVA with Dunnett’s post-hoc. ns, not significant. **p<0.01, ***p<0.001, ****p<0.0001.

Using our high-throughput automated curling assay, we were able to test a total of 400 conditions to study disease mechanisms^11^, as well as to screen for drugs that improve motor function in *bcat-1(RNAi)* worms (Figure 1h, 2b). More than 32,000 worms were tested, with an average of 83 animals per condition and 13 worms per well. 19% of the worms could not be analyzed because they became entangled/overlapping during the assay and were therefore automatically censored. The remaining 81% represent ~1,050,000 detected objects across all the captured snapshots, with 63% of these having passed as “successfully found” individual worms after 11% computer and 6% manual censoring (Figure 1h). On average, ~2100 images of successfully detected worms were used to quantify curling for each experimental condition.

To identify new potential treatments for PD, we conducted a proof-of-concept screen of 50 FDA-approved drugs for those that improve *bcat-1*-related motor dysfunction in *C. elegans*. Neuronal RNAi-sensitive worms were fed *bcat-1* or control RNAi at the onset of adulthood until day 4, and were then switched to heat-killed OP50 *E. Coli* bacteria with 50 µM drug or vehicle (0.5% DMSO). Curling assays were performed on day 8 (Figure 1a). We specifically chose a mid-stage intervention (day 4) in order to mimic the PD patient who enters the clinic seeking treatment at a time when the disease is already underway.

The 50 drugs in the screen were selected to represent a wide range of clinical indications (Supplementary Figure 1a) and targets/mechanisms of action (Supplementary Figure 1b). Half of the drugs are known to target neurotransmission, including medications for epilepsy, depression and other mood disorders, schizophrenia, and the neurodegenerative diseases Alzheimer’s and amyotrophic lateral sclerosis. Drugs with non-neurological indications were included to allow for the discovery of highly novel therapies; current indications for these drugs include cancers, diabetes, infectious diseases, autoimmune disorders, and cardiovascular diseases.

Several drugs emerged from the screen as potential candidates for repurposing to PD. Metformin, enasidenib, nitisinone, felodipine, and ethosuximide reduced curling to <60% of vehicle-treated levels (Figure 2c). Post-screening validation with automated analysis of 30s videos confirmed that 4 out of 5 of these drugs reliably and significantly reduced curling (Figure 2d), and final confirmation for our top candidate, metformin, was obtained by manual hand-counting (Figure 2e). Detailed characterization of metformin’s neuroprotective action in *bcat-1(RNAi)* worms is reported in our companion study^11^.

While only one of the four validated drugs from our screen, ethosuximide, is currently prescribed for a neurological disease, the known targets/activities of the four identified candidates appear to converge on similar cellular pathways related to metabolism (Figure 2f). Metformin and nitisinone act on metabolic targets^12, 13^, enasidenib inhibits the TCA cycle enzyme IDH-2^14^, and ethosuximide is a calcium channel blocker^15^. These data suggest that drugs with metabolic and/or mitochondrial targets may represent a new class of therapies for PD.

*C. elegans* is highly amenable to high-throughput screening approaches, allowing for rapid testing of potential disease treatment options^16^. Combined with our user-friendly program that facilitates intuitive analysis of recorded snapshots, our high-throughput behavioral testing platform can quickly screen curling behavior across thousands of worms. We have screened more than 32,000 worms at a rate 20 times faster than the alternative manual assay, focusing on uncovering the mechanisms by which reduction of *bcat-1* is neurotoxic^11^ and testing for potential compounds that may be repurposed for PD. We identified the FDA-approved drugs metformin, enasidenib, nitisinone, and ethosuximide as highly promising candidates for further investigation as potential late-in-life interventions in PD. Furthermore, the application of our high-throughput assay is not limited to worms with *bcat-1* knockdown; it can also be used to study straight, bent, and coiled swimming postures with other RNAi treatments or in other genetic strains (such as cholinergic mutants *cha-1* and *unc-17*; see Figure 2g). Collectively, these findings point to the utility of our high-throughput platform for uncovering disease mechanisms and potential therapeutics.

## Methods

### *C. elegans* Strains and Maintenance

*C. elegans* strains were grown at 20°C on nematode growth medium (NGM) plates or high growth medium (HG) plates seeded with OP50 *Escherichia coli* or HT115 RNAi *Escherichia coli*. RNAi clones were obtained from the Ahringer RNAi library. The following strains were used in this study: wild-type worms of the N2 Bristol strain, CQ511 [*sid-1(pk3321)*]; uIs69 [pCFJ90 (*myo-2p::mCherry, unc-119p::sid-1*)], CF512 (*fem-1(hc17); fer-15(b26)*), CQ434 baIn11 [*dat-1p::αsynuclein; dat-1p::gfp*]; vIs69 [pCFJ90 (*myo-2p::mCherry + unc-119p::sid-1*)], CQ491 vIs48 [*unc-17p::gfp*]; vIs69 [pCFJ90 (*myo-2p::mCherry + unc-119p::sid-1*)], TU3311 uIs60 (*unc-119p::sid-1, unc-119p::yfp*), CB113 (*unc-17(e113*)), TY1652 (*cha-1*(*y226*)).

### Drug treatments

For RNAi experiments, worms were synchronized from eggs by bleaching and placed on HG plates seeded with OP50. For CF512 (*fem-1(hc17); fer-15(b26)*), animals were sterilized by incubation at 25°C from L2-L4. RNAi-seeded 100-mm NGM plates containing carbenicillin and IPTG were pre-induced with 0.1 M IPTG 1h prior to transfer of worms at the L4 stage. In all experiments, control RNAi refers to empty vector pL4440 in HT115 *Escherichia coli*. Day 4 worms were transferred onto fresh NGM plates seeded with 1 mL heat-killed OP50 bacteria and 50-µM drug or vehicle (0.5% DMSO). OP50 was killed by incubation at 65°C for 30 min. Curling was measured on day 8. Drugs for high-throughput screening were selected from the FDA-approved Drug Library (MedChem Express).

### Manual assay

10-15 worms at a time were picked into a 10 µL drop of M9 buffer on a microscope slide^5^. Approximately eight to ten 30s videos were captured for each tested condition using an ocular-fitted iPhone camera attached to a standard dissection microscope and curling was quantified with a standard EXTECH Instruments stopwatch. % time spent curling is defined as the sum of the periods in which either the head or tail makes contact with a noncontiguous segment of the body, divided by the total time measured.

### Automated image capturing

On the day of analysis, animals were washed twice with M9 buffer and dispensed into 35mm petri dishes rocking on an orbital shaker to ensure even distribution. Worms were pipetted with large orifice tips into the wells of a 96-well plate. 30s videos or a series of snapshots were obtained for individual wells containing up to 30 worms each. It is imperative to pre-fill 35mm petri dishes and the 96-well plate with 6-mg/ml OP50 solution in M9 to prevent starvation and worms sticking to surfaces. Before initiating each round of automated image capturing, worms in the 96-well plate were stimulated on a thermoshaker (Eppendorf) at 900rpm for 60s. Worms are most active in the first 2 hours after being transferred into liquid buffer; worm movement gradually decreases over time^17^. However, up to five rounds of automated snapshot recordings, seven snapshots per round, can be completed in less than one hour. Images were captured with inverted colors, i.e., white worms on a black background. To ensure efficient image capturing, rows were sequentially imaged in alternating directions.

### Curling quantification software

After importing raw images, the user roughly determines the circular outline of one well in the first snapshot by identifying 3 points on the well’s circumference. To locate the animals in a single image, the gradient of image delineates the edges of the animals. Detection parameters such as min/max area-limit, Mask-area, Levels-Otsu’s method, N-node, and Jaggedness can be adjusted based on imaging setup or user preference. Detected objects are initially filtered through a minimum and maximum target surface area. The defaults are 0.1 and 8 times the average size of an adult worm. Mask-area determines the size of the mask around found objects. Otsu’s levels can accept combination values such as [1,1], [2,1], [2,2], etc., where the first number identifies how many multilevel thresholds will be calculated for the gradient image using Otsu’s method, and the second number dictates which threshold will be used for initial binarization. The default values are set to [1,1]. N-node indicates the number of nodes used to outline each detected object with default value of 60. Jaggedness threshold automatically filters out chunks of bacteria when found objects are not sufficiently smooth. Jaggedness is defined as the perimeter of the originally found object to the perimeter of the smoothened one; the default value is 1.3.

For the second round of filtering, Otsu’s method is used to identify area and circumference outliers when analyzing the found objects. The first number in Area-filter and Perimeter-filter indicates the number of multilevel thresholds used. Then, objects that have smaller or larger area/perimeter are censored. Zero denotes no high or low band filtering. Additionally, Dim-filter removes extremely thin or long objects.

Utilizing circularity and convex hull shape factors, the program categorizes animals into curled, near-curled, or non-curled subgroups. Circularity is defined as the degree to which the found object is similar to a circle. Circularity is a measure of both form and roughness. Outlining all found objects with the same number of nodes, circularity will only represent form. The further away from a circle, the lower the circularity value. Near-curled and non-curled worms have circularity values of 0.1-0.3. A cut-off value of 0.45 is used to identify curled worms (defined as either the head or tail making contact with a noncontiguous segment of the body). To identify near-curled worms, a new boundary enclosing perimeter nodes of the found object is redrawn using a shrink factor between 0 and 1. Zero results in the convex hull while 1 gives the compact boundary enveloping the original perimeter nodes. If the circularity of the redrawn outline is significantly bigger than the circularity of the original boundary (circularity ratio), worms are categorized as near-curled. Default values for shrink factor and circularity ratio are 0.7 and 2, respectively.

The user can review color-coded worms in a montage image to manually override automated decisions. While left-click toggles between curled, censored, and non-curled labels, right-click toggles between near-curled, censored, and non-curled labels. Using this high-throughput workflow, three rounds of snapshots with a total of 21 snapshots per well is sufficient for curling analysis.

### Statistical analysis

For all comparisons between two groups, an unpaired Student’s t-test was performed. For comparisons between multiple groups, One-Way ANOVA was performed with post-hoc testing. GraphPad Prism was used for all statistical analyses.

## Acknowledgements

We thank the *C. elegans* Genetics Center for strains (P40 OD010440); and the Murphy lab for discussion. This work was supported by the Glenn Foundation for Medical Research award to C.T.M. (GMFR CNV1001899) and NIH awards (grant #1RF1AG057341 (NIA), 5R01AG034446 (NIA), and 5DP1GM119167 (NIGMS)). C.T.M. is the Director of the Glenn Center for Aging Research at Princeton and an HHMI-Simons Faculty Scholar. D.E.M. was supported by Ruth L. Kirschstein NRSA (NIA F32AG062036).

## Author contributions

S.S., D.E.M., and C.T.M. designed experiments. S.S., D.E.M., R.K. and W.K. performed experiments and analyzed data. S.S., D.E.M., and C.T.M wrote the manuscript.

---

**Supplementary Figure 1.**
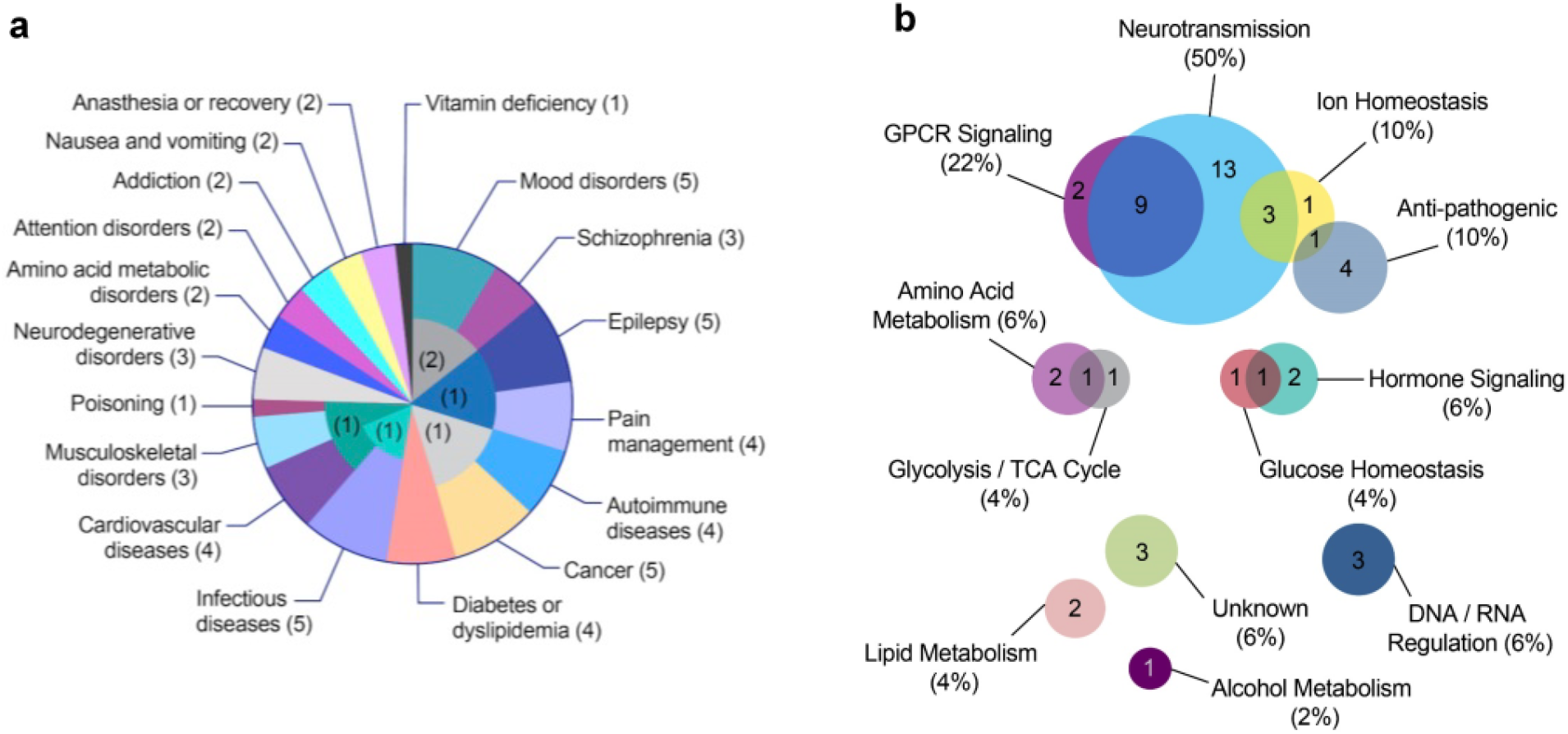
(a) Disease indications of the 50 drugs selected for the screen, with number of drugs in parentheses. (b) Targets/ mechanisms of action of the 50 drugs, with percentage of drugs in parentheses.

